# kegg_pull: a Software Package for the RESTful Access and Pulling from The Kyoto Encyclopedia of Gene and Genomes

**DOI:** 10.1101/2022.11.03.515120

**Authors:** Erik Huckvale, Hunter N.B. Moseley

## Abstract

**Background:** The Kyoto Encyclopedia of Genes and Genomes (KEGG) provides organized genomic, biomolecular, and metabolic information and knowledge that is reasonably current and highly useful for a wide range of analyses and modeling. KEGG follows the principles of data stewardship to be findable, accessible, interoperable, and reusable (FAIR) by providing RESTful access to their database entries via their web-accessible KEGG API. However, the overall FAIRness of KEGG is often limited by the library and software package support available in a given programming language. While R library support for KEGG is fairly strong, Python library support has been lacking. Moreover, there is no software that provides extensive command line level support for KEGG access and utilization.

**Results:** We present kegg_pull, a package implemented in the Python programming language that provides better KEGG access and utilization functionality than previous libraries and software packages. Not only does kegg_pull include an application programming interface (API) for Python programming, it also provides a command line interface (CLI) that enables language agnostic utilization of KEGG for a wide range of data analysis pipeline use-cases. As kegg_pull’s name implies, both the API and CLI provide versatile options for pulling (downloading and saving) an arbitrary (user defined) number of database entries from the KEGG API. Moreover, this functionality is implemented to efficiently utilize multiple central processing unit cores. Many options are provided to optimize fault-tolerant performance across a single or multiple processes, with recommendations provided based on extensive testing and practical network considerations.

**Conclusions:** The new kegg_pull package enables new flexible KEGG retrieval use cases not available in previous software packages. The most notable new feature that kegg_pull provides is its ability to robustly pull an arbitrary number of KEGG entries with a single API method or CLI command, including pulling an entire KEGG database. We provide recommendations to users for the most effective use of kegg_pull according to their network and computational circumstances.

## Background

The Kyoto Encyclopedia of Genes and Genomes (KEGG) [1][2][3] is a collection of databases containing organized biomolecular and metabolic data (information) for over 3000 species with sequenced genomes. A primary component of each KEGG database is a KEGG entry, a relational table record that represents and describes a specific chemical, biochemical, or biological entity (e.g. a chemical compound, a biochemical reaction or pathway, an enzyme, a gene, a species etc.). Each KEGG entry is uniquely identified with a KEGG ID. The KEGG databases are updated regularly and made publicly available via the KEGG website [4]. However, the website is designed for manual access through a web browser. For more automated access, KEGG provides a Representational State Transfer (REST) web application programming interface (web API). A REST web API is a predominant software architecture for making uniform interactions between software components via the world wide web. These interactions typically occur as requests in the form of a uniform resource locator (URL) provided through the http protocol, with a “GET” http request fetching data from a web server and a “PUT” http request typically updating data on a web server [5]. The KEGG REST web API (KEGG API) [1] provides a set of operations for accessing most of the organized data in KEGG as described on the KEGG API web page: https://www.kegg.jp/kegg/rest/keggapi.html. In particular, the KEGG API enables researchers to retrieve KEGG data, especially KEGG entries, for use in their own analyses. Operations to obtain KEGG entry IDs include the “list” and “find” operations, the output of these operations returning meta data which needs to be parsed out if only the entry IDs themselves are desired. And the “get” operation provides KEGG entries themselves given their corresponding IDs.

Users can make requests to REST web APIs by providing the correct URL to a variety of web accessing software, for example a web browser, library packages like the Python requests module [6], and even command line tools like cURL [7]. However, construction of these URLs is somewhat cumbersome, requiring specific URL templates for a specific REST web API with some URL construction expertise, which is even limiting for some bioinformaticians, let alone biologists with limited computational skills. Library packages do exist both in R [8] and Python [9] for accessing most of the KEGG API. However, to our knowledge, none of these packages provide a command line interface (CLI) for researchers who cannot or prefer not to write code. Also missing is a package that provides a variety of other use cases, for example obtaining KEGG entry IDs alone with the metadata already parsed out or downloading an arbitrary number of entries in a single command. Therefore, we introduce a new Python package kegg_pull, which meets the above use cases and more. We have implemented kegg_pull to a rigorous industrial standard, which includes both unit and integration tests. The kegg_pull package is installable through the Python Package Index (https://pypi.org/project/kegg-pull/).

We created kegg_pull to promote the FAIR (Findable, Accessible, Interoperable, and Reusable) guiding principles of data stewardship [10] with respect to KEGG. While KEGG is primarily responsible for implementing FAIR, kegg_pull improves on the accessibility, interoperability, and reusability of the KEGG API. The kegg_pull package improves the accessibility by making the utilities of the KEGG API accessible to Python programmers, including those that may have limited knowledge of web development. Additionally, it makes these utilities accessible to command line users. Interoperability is improved by making the output from the KEGG API available in a form suitable for other contexts, such as Python objects in a python script, files in the file system, and console output that can be piped into another command on the command line. This necessarily allows KEGG data to be used in shell scripts. The improved interoperability enables the output to be transferred downstream within a complex workflow or to be used by a workflow manager. Finally, the kegg_pull package improves reusability by making KEGG data more easily reused by researchers in a variety of Python-based data analyses and command line-based data analysis pipelines.

### Implementation

The kegg_pull package provides several useful CLI and API features for interacting with the KEGG API. This includes wrapper methods/commands for all the REST API operations, pulling lists of KEGG entry IDs, and pulling an arbitrary number of KEGG entries that are automatically separated and saved into individual files, all with a single function call or command line execution. The kegg_pull API is implemented in four submodules (Figure 1): pull, entry_ids, rest, and kegg_url. The kegg_pull CLI reuses this API to provide a higher level of functionality, conveniently accessible from the command line without needing to write Python scripts. If more flexibility is necessary, however, researchers with programming expertise can use the kegg_pull API in their own Python scripts and programs.

**Figure 1.**
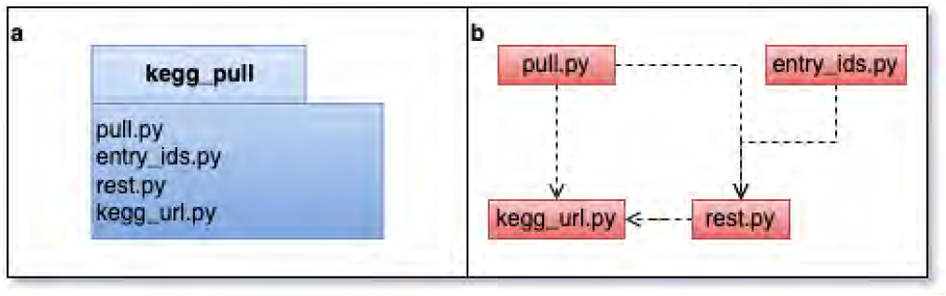
**a.** UML package diagram. **b.** Submodule dependencies.

The kegg_url submodule constructs URL objects for accessing the KEGG REST API. As illustrated in Figure 2, each URL class inherits from the AbstractKEGGurl class with the shared “url” property, which is the constructed URL string itself. Each inheriting concrete class corresponds to one of the KEGG REST operations. Some of the KEGG REST operations have two forms, necessitating two different URL classes to represent both forms.

**Figure 2.**
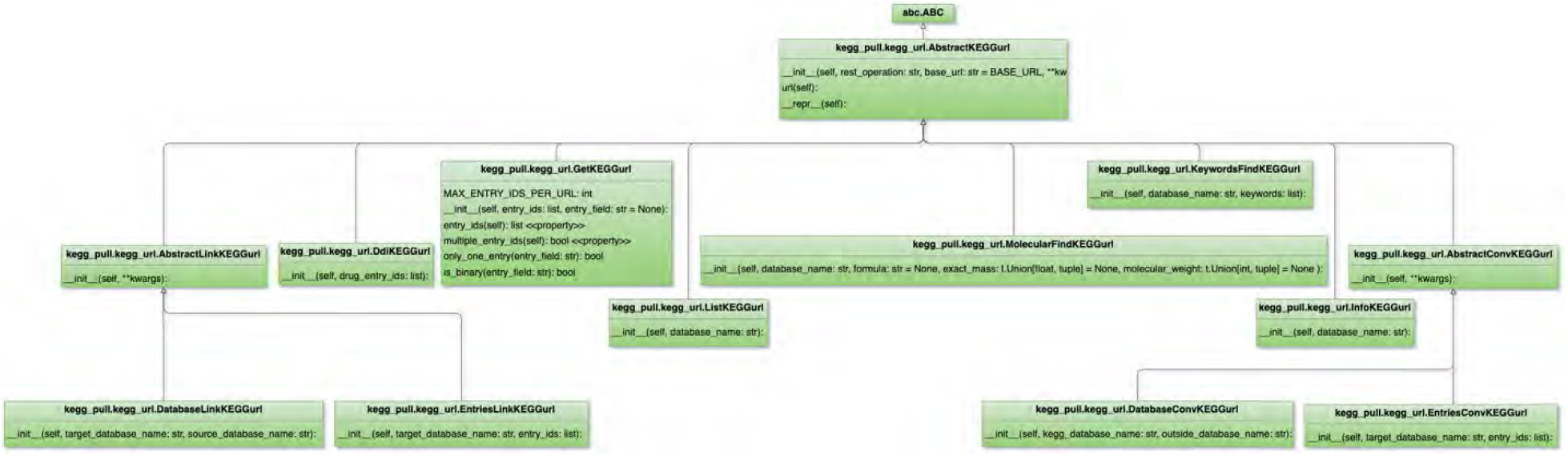
Class diagram of the 6eg_url.py submodule.

The rest submodule depends on the kegg_url submodule for providing wrapper methods over the KEGG REST API operations, contained within the KEGGrest class (Figure 3). Each wrapper method uses its corresponding KEGG URL class. A user-created Python program could use the kegg_url submodule to construct the URLs and, if more control over the URLs is needed, pass them into a Python library such as requests. However, the benefits of using the wrapper methods of the KEGGrest class include:

1. Abstracting the URL strings so less knowledge of web development is needed and using the requests library under the hood automatically.
2. Allowing the caller to specify the number of tries to make a request in case initial requests fail or time out.
3. Allowing the user to specify how long requests should wait for a response before being marked as timed out.
4. Allowing the caller to specify the sleep time in between requests that time out or are blacklisted to give the KEGG web server time to return to an accessible state. Blacklisting is when the KEGG web server temporarily blocks further requests when it deems too many have been made, necessitating waiting until the blacklisting is repealed.
5. Returning a KEGGresponse object (see Figure 3) which contains the information from a response generated from a request to the KEGG API, including both a text body and binary body if applicable, the URL constructed for the request, and the status (i.e. SUCCESS, FAILED, or TIMEOUT).

**Figure 3.**
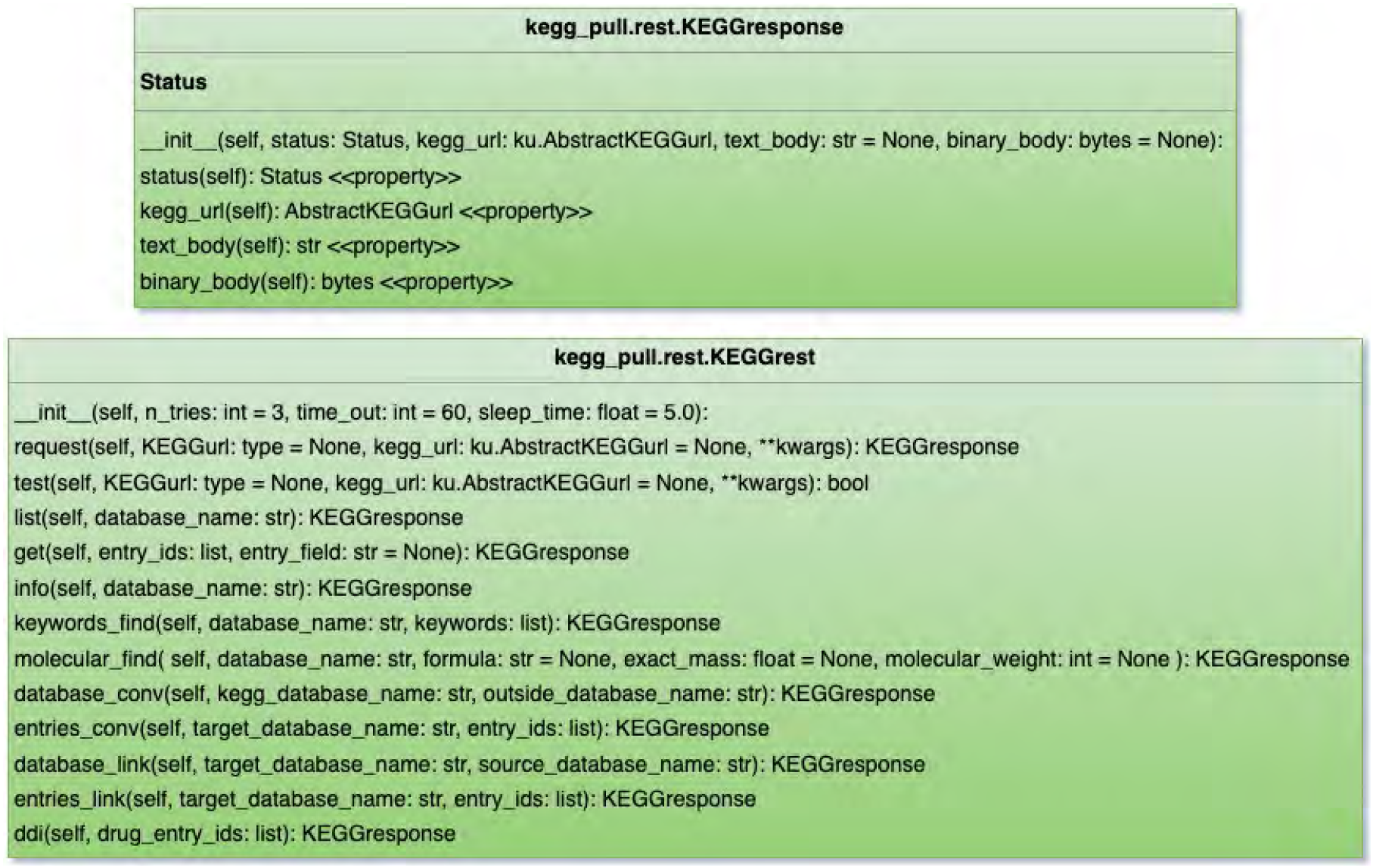
Class diagram of the rest.py submodule.

The entry_ids submodule depends on the rest submodule, since it uses the KEGGrest class’s methods for obtaining entry IDs (i.e. the “list” method for getting all the entry IDs of a given KEGG database and the keywords_find and molecular_find methods for getting the entry IDs using the “find” KEGG REST operation.). These methods are contained within the EntryIdsGetter class which contains a KEGGrest object, allowing the user to customize how requests are made if desired (e.g. the time to wait for a time out, etc.). A user-created Python program could use the KEGGrest class to directly get the entry IDs from its relevant methods, but the response body comes in a string that contains metadata on top of the entry IDs. The EntryIdsGetter class will additionally parse the string to return a list containing only the entry IDs themselves. The EntryIdsGetter class (Figure 4) additionally contains a method for loading a list of entry IDs from a file if the user already has the entry IDs they’d like to retrieve in their local file system.

**Figure 4.**
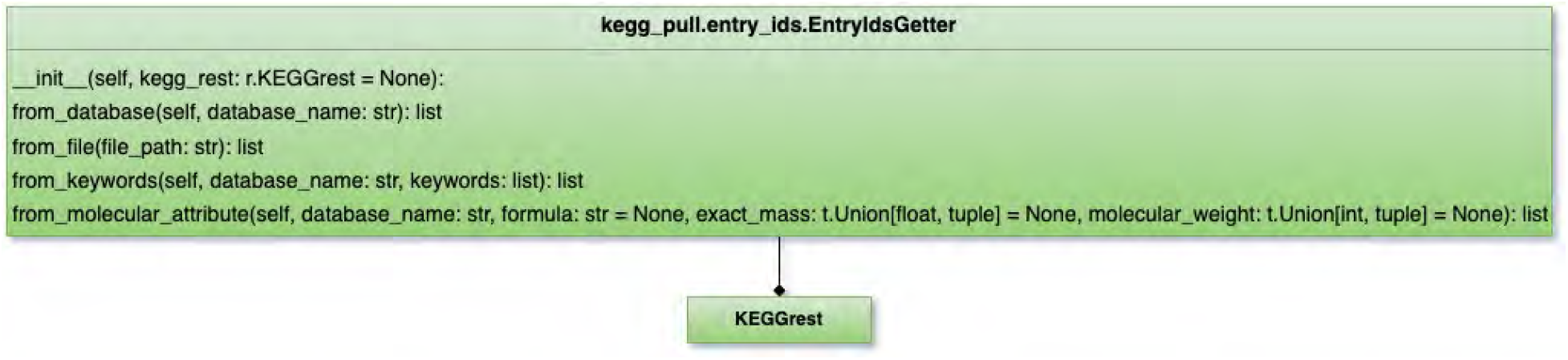
Class diagram of the entry_ids.py submodule.

The pull submodule depends on the rest submodule for obtaining KEGG entries given their IDs. Within the submodule is the SinglePull class which contains a KEGGrest object to use its “get” method followed by extracting one or more KEGG entries from the text body (string) of the KEGGresponse returned or the binary body (bytes) in the case of “image” fields from relevant entries. While “image” can be set as the entry_field parameter to the pull method, all the other entry fields for the “get” KEGG REST operation can be set with this parameter to pull particular fields from entries. If unspecified, the entry itself will be pulled in its standard format. The SinglePull class (Figure 5) performs just one request in its pull method, as its name suggests. Currently, there is a limit of 10 entry IDs that can be sent in this request, since KEGG API’s response will truncate to 10 entries if more are requested. So, the SinglePull class is limited on how many entries it can pull for a single command. One could use the KEGGrest class’s get method directly to obtain entries from the response body. However, the benefits of using the pull method of the SinglePull class are:

1. The string or bytes response is automatically saved to the file system with the entry ID as the file name and the entry field as the extension, the “.txt” extension used if no entry field is specified.
2. If the response body is binary, the file is automatically saved in binary format.
3. If multiple entry IDs are provided, the entries are automatically split by their respective delimiter in the response body and saved separately in individual files, sparing the user from needing to perform the same empirical experiments we did during software development to determine what the delimiters are in the first place and additionally sparing them from needing to write their own parser functions.
4. If multiple entries are requested and the initial request fails or not all requested entries were returned, each entry is requested one at a time (instead of them all being requested in a single response) to maximize the number of successful entries pulled.
5. The user can specify to save the output file in a regular directory or a zip archive file. If the provided directory name ends in “.zip”, the file is automatically saved in a ZIP archive of that name. If either the provided directory or provided ZIP archive doesn’t already exist, one will be automatically created.
6. A PullResult object (Figure 5) is returned specifying by their ID which of the entries requested were successfully pulled, which entries failed to be pulled, and which entries timed out.

**Figure 5.**
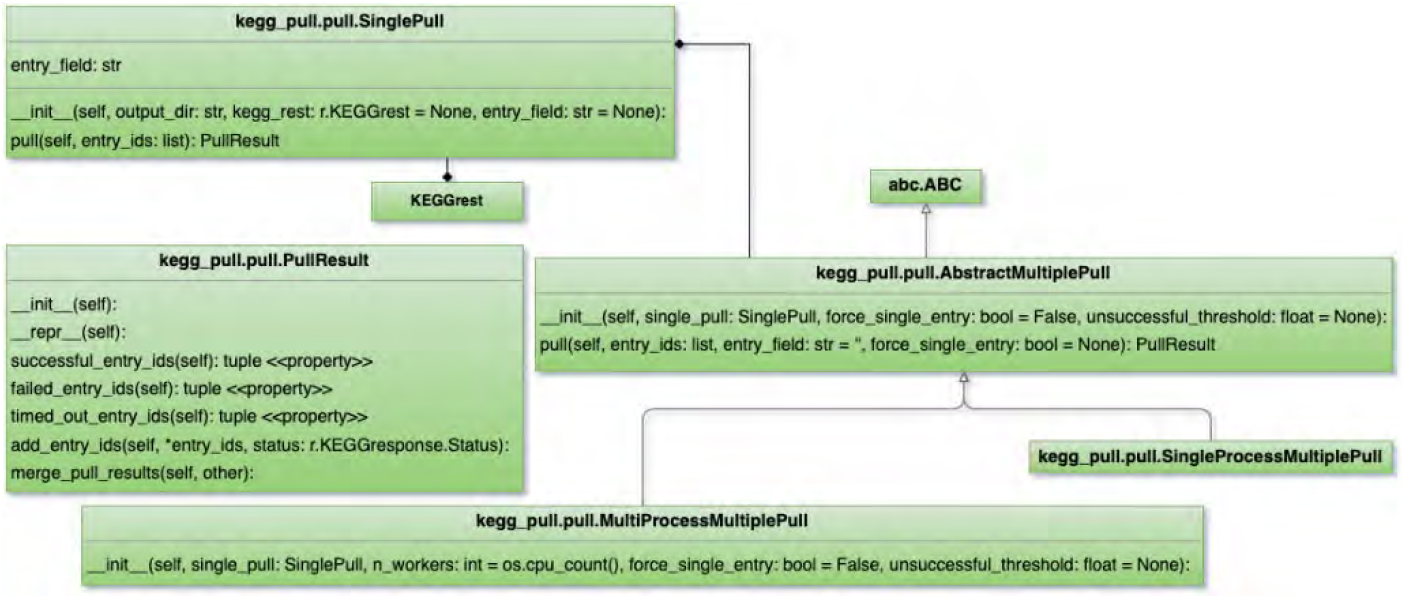
Class diagram of the pull.py module.

The pull submodule provides additional classes for pulling an arbitrary number of entry IDs, namely SingleProcessMultiplePull and MultiProcessMultiplePull (see Figure 5). These contain a SinglePull object and call its pull method as many times as necessary to request all the entry IDs provided to their own pull method. As a result, there is no limit to the number of entry IDs unlike with the SinglePull class. The pull method of these classes, like that of the SinglePull class, also returns a PullResult object detailing which of the requested entry IDs succeeded, failed, or timed out. A user-created Python program could have its own loop which calls a SinglePull object’s pull method multiple times if desired. However, the SingleProcessMultiplePull and MultiProcessMultiplePull objects already have this functionality along with merging the individual PullResult return values of the individual SinglePull calls into a comprehensive PullResult object. There is also a progress bar displayed in the console and the ability to halt the program if too many of the requests fail or time out. The user can also specify a failure rate threshold for automatic halting.

In addition, the MultiProcessMultiplePull class enables pulling entries across multiple processes to pull more entries in less time when running on a system with multiple cores. The user can specify the number of processes to use, the default being the number of cores available. While multiprocessing is safe in the case of saving files to a regular directory since each file is written entirely within its own process rather than multiple processes writing to that same file, it is not safe when writing files to a ZIP archive. While the processes are writing different files to this ZIP archive, the ZIP archive itself is technically a single file which multiple processes write to. Having multiple processes writing to a single ZIP archive creates a race condition, which will corrupt the ZIP archive when multiple processes open and write to it at the same time. Therefore, the MultiProcessMultiplePull object uses a multiprocessing.Lock object stored in its SinglePull object to process lock the code block that writes to the ZIP archive, meaning while one process opens and writes to the ZIP archive, other processes are temporarily blocked from doing so. This means a slight reduction in performance since the other processes are delayed whenever they happen to come to the ZIP archive writing code at the same time as another process, but the majority of the code executed across multiple processes is not locked, resulting in an increase in efficiency even when writing to a ZIP archive file (See table 5).

**Table 1.** Pull Success Percentage And Time Spent Pulling By Sleep Time - KO Database (25,439 entries)

| **Sleep Time**<br>**(seconds)** | **0.0** | **0.5** | **1.0** | **2.0** | **3.0** | **5.0** | **10.0** |
| --- | --- | --- | --- | --- | --- | --- | --- |
| **Percent Success** | 94.68 | 94.89 | 96.78 | 99.78 | 100.0 | 100.0 | 100.0 |
| **Pull Time**<br>**(minutes)** | 12.99 | 16.03 | 14.69 | 10.82 | 8.51 | 8.44 | 8.7 |
| *Number of minutes spent attempting to pull all the entries in the KO KEGG database. Percent success is the percentage of the entries in the KO database that were successfully pulled while the others failed. Difference in pull time and percent success varies by the --sleep-time option on the kegg_pull CLI. All other options remained the same, including the use of multiprocessing. Values were collected on a 12 core (hyperthreaded) machine using 12 processes.* |  |  |  |  |  |  |  |

**Table 2.**
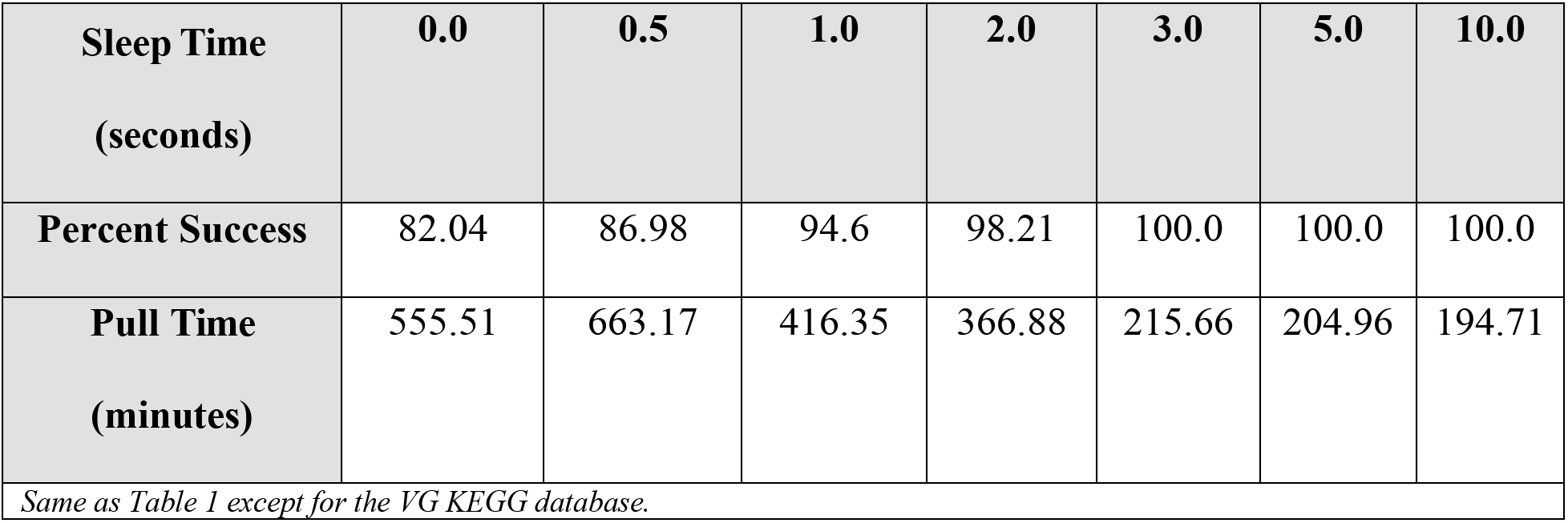
Pull Success Percentage And Time Spent Pulling By Sleep Time - VG Database (595,443 entries)

| **Sleep Time**<br>**(seconds)** | **0.0** | **0.5** | **1.0** | **2.0** | **3.0** | **5.0** | **10.0** |
| --- | --- | --- | --- | --- | --- | --- | --- |
| **Percent Success** | 82.04 | 86.98 | 94.6 | 98.21 | 100.0 | 100.0 | 100.0 |
| **Pull Time**<br>**(minutes)** | 555.51 | 663.17 | 416.35 | 366.88 | 215.66 | 204.96 | 194.71 |
| *Same as Table 1 except for the VG KEGG database.* |  |  |  |  |  |  |  |

**Table 3.**
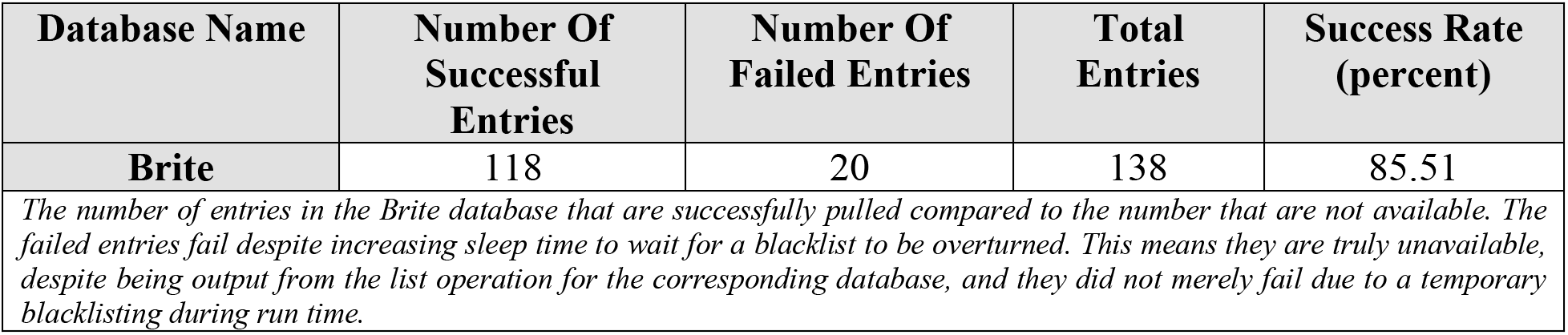
Failed Entries In The Brite Database Regardless Of Sleep Time

**Table 4.** Multi-process Pull Time Vs. Single Process Pull Time (minutes) Into A Regular Directory

| Database Name | Multi-process Pull Time | Single Process Pull Time | Number Of Entries |
| --- | --- | --- | --- |
| **Pathway** | 0.1 | 1.05 | 558 |
| **Compound** | 6.4 | 73.62 | 19,004 |
| **KO** | 8.32 | 74.0 | 25,458 |
| *The amount of time to pull and save all the entries of a given database on a single process (one core) compared to pulling across multiple processes (multiple cores). The above values result from running kegg_pull on a 12 hyper-threaded core machine using 12 processes for multiprocessing and one process for single-processing. The sleep time and all other options for each were also constant. Files were saved in a regular directory.* |  |  |  |

**Table 5.** Multi-process Pull Time Vs. Single Process Pull Time (minutes) Into A ZIP Archive

| Database Name | Multi-process Pull Time | Single Process Pull Time | Number Of Entries |
| --- | --- | --- | --- |
| Pathway | 0.42 | 1.13 | 558 |
| Compound | 38.09 | 73.62 | 19,004 |
| KO | 66.74 | 138.44 | 25,458 |

Same as table 4 except files were saved in a ZIP archive.

The top-level command line interface usage description in Figure 6 shows that kegg_pull has 3 subcommands, namely rest, entry-ids, and pull. These subcommands reuse the rest, entry_ids, and pull submodules and are analogous to the entities within them. However, the command line interface provides additional functionality. For the rest and entry-ids subcommands, the user can choose whether to print the output to the console or save it in a file. Similar to the pull methods in the API, the user can choose to save within a regular directory or within a ZIP archive. Using the pull subcommand on the command line makes the progress bar visible in the console and saves the information contained within the PullResult to a file. Not only are the successful, failed, and timed out entries specified (by their ID) in this file, but other useful information about the pull is saved as well, including the time it took to pull all the requested entries, the success percent or percent of entries that succeeded out of the total number of requested entries, and the amount of each entry ID category, i.e. the number of entries that succeeded to be pulled, the number that failed, etc. If the user instructs kegg_pull to abort upon too many entries not succeeding, a file detailing the results of the aborted pull is created.

**Figure 6.**
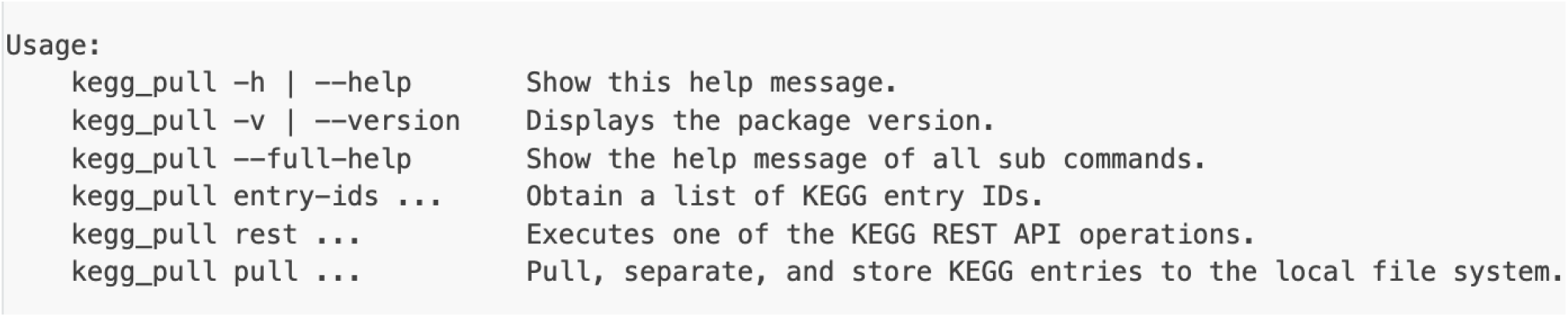
Top Level Command Line Usage of kegg_pull

More details on the kegg_pull API and CLI is available in the online package documentation: https://moseleybioinformaticslab.github.io/kegg_pull/.

## Results

The kegg_pull CLI enables the user to pull all the entries in a specified KEGG database with a single command. We discovered that the time it takes to accomplish this varies based on the – sleep-time option (the time to wait in between timed out requests and blacklisted requests). This option also affects the success percentage, the percentage of entries that succeed rather than fail. When we performed the execution time experiments (Tables 1 and 2), we found that none of the requests timed out, so the results most likely reflect the percentage of successfully pulled entries as compared to those that were blacklisted for all three tries. Since each request only tried 3 times, waiting for 0 seconds in between tries would not give enough time to wait for the KEGG web server to repeal the blacklisting. This is most likely why we see an increase in the success percentage as the sleep time increases. After reaching 100 percent, increasing the sleep time unsurprisingly no longer affects the success percentage. Our results in Table 1 also show a negligible increase in pull time after increasing sleep time past reaching 100 percent success in the case of the KO database. In the case of the larger VG database shown in Table 2, we actually see a continued decrease in pull time after reaching 100 percent success. See the supplementary materials for the single process version of this experiment (ko database only). From that table, we see that even a sleep time of 0.0 can result in 100 percent success when pulling in a single process.

In the case of the Brite KEGG database, 20 entries consistently failed despite increases in sleep time. We can conclude that these 20 entries are simply unavailable rather than resulting from indeterministic blacklisting. The “list” KEGG REST operation provides the entry IDs of an entire KEGG database. After attempting to pull the entries corresponding to the Brite IDs returned by the “list” operation, not all of the entries were available as tabulated in Table 3. See the supplementary materials for the list of the Brite entry IDs that fail.

When making multiple requests to the KEGG REST API to pull an arbitrary number of entries, a kegg_pull user can specify in both the API and CLI to use one process or multi-processing. As illustrated in Tables 4 and 5, we see that the pull time for whole KEGG databases can be dramatically reduced when using multi-processing.

We see that pulling KEGG entries into a ZIP archive significantly increases pull time as compared to pulling into a regular directory. However, multi-process pulling into a ZIP archive is still substantially faster than single process pulling into a ZIP archive, despite process locking the code block that accesses the ZIP file, which is required to prevent corrupting the ZIP archive file.

Table 6 demonstrates the substantial increase in pull efficiency from the KEGG API’s ability to request multiple entries within a single response body. The success percentage can also decrease slightly when only pulling one entry per request, necessitating increased sleep time.

**Table 6.** Pull Time (minutes) And Percent Success With One Entry At A Time Vs. Ten Entries At A Time And Different Sleep Times

| Database Name | One Entry At A Time |  |  |  | Ten Entries At A Time |  |  |  |
| --- | --- | --- | --- | --- | --- | --- | --- | --- |
|  | Sleep Time 5 Seconds |  | Sleep Time 20 Seconds |  | Sleep Time 5 Seconds |  | Sleep Time 20 Seconds |  |
|  | Percent Success | Pull Time | Percent Success | Pull Time | Percent Success | Pull Time | Percent Success | Pull Time |
| Module | 98.69 | 1.34 | 100.0 | 1.59 | 100.0 | 0.06 | 100.0 | 0.07 |
| Pathway | 98.75 | 1.6 | 100.0 | 1.67 | 100.0 | 0.11 | 100.0 | 0.09 |
| Compound | 99.24 | 63.62 | 100.0 | 61.36 | 100.0 | 6.77 | 100.0 | 7.21 |
| KO | 99.39 | 83.91 | 100.0 | 90.4 | 100.0 | 8.42 | 100.0 | 9.22 |

The amount of time (minutes) to pull all the entries from a given database and the success percentage when pulling one entry at a time (with the --force-single-entry flag set) compared to pulling ten entries (maximum allowed by KEGG) per request. Each of these are compared to a lower sleep time vs. a higher sleep time. Results were collected on a 12 (hyperthreaded) core machine on 12 processes with all other options consistent.

When pulling entries from KEGG, there is a maximum number of entries that can be pulled in a single request due to KEGG API response limitations. While the entries of all the KEGG databases, except for Brite, support requesting this maximum amount, that is not necessarily the case when a user desires to pull particular fields from the entries. For example, a user might want to pull an amino acid sequence from a gene entry or a mol file from a compound entry. Some entry fields allow this maximum number while others do not. While this is not currently specified in the KEGG REST documentation, we empirically discovered which entry fields allow this and which only allow a single entry to be pulled at a time. With this information shown in Table 7, we implemented kegg_pull such that it will pull only one entry at a time if the user wants an entry field that does not support multiple for a single request. Likewise, if the user specifies to pull all the entries from the Brite database, kegg_pull will only pull one entry at a time in that case as well. That convenience for the Brite database, however, is only available in the CLI when the database name is specified. When pulling Brite entries in all other cases, there isn’t a way to tell which database the entries are coming from, necessitating the force_single_entry parameter for the API and the – force-single-entry option for the CLI. Even if the user neglects to set this parameter / option, however, kegg_pull is smart enough to retry on each requested entry individually if not all of the requested entries are pulled initially. Forcing a single entry at a time is for efficiency rather than successful pulls.

**Table 7.** Entry Fields That Allow Multiple Entries To Be Pulled Vs. Those That Only Allow One Per Request

| Entry Field | Can Pull Multiple Entries In One Request |
| --- | --- |
| aaseq | ✓ |
| ntseq | ✓ |
| mol | ✓ |
| kcf | ✓ |
| image | ✗ |
| conf | ✗ |
| kgml | ✗ |
| json | ✗ |

Since the CLI builds off of the API, a kegg_pull user can write API code that’s analogous to corresponding CLI commands. We say analogous rather than synonymous because the CLI can do more than the analogous API commands (e.g. saving the output to a file or printing to standard output rather than merely returning a Python object). When a user chooses to use the API over the CLI, they sacrifice potential convenience for higher control, if needed. Table 8 has examples of prominent API usage followed by their analogous CLI commands in Table 9.

**Table 8.**
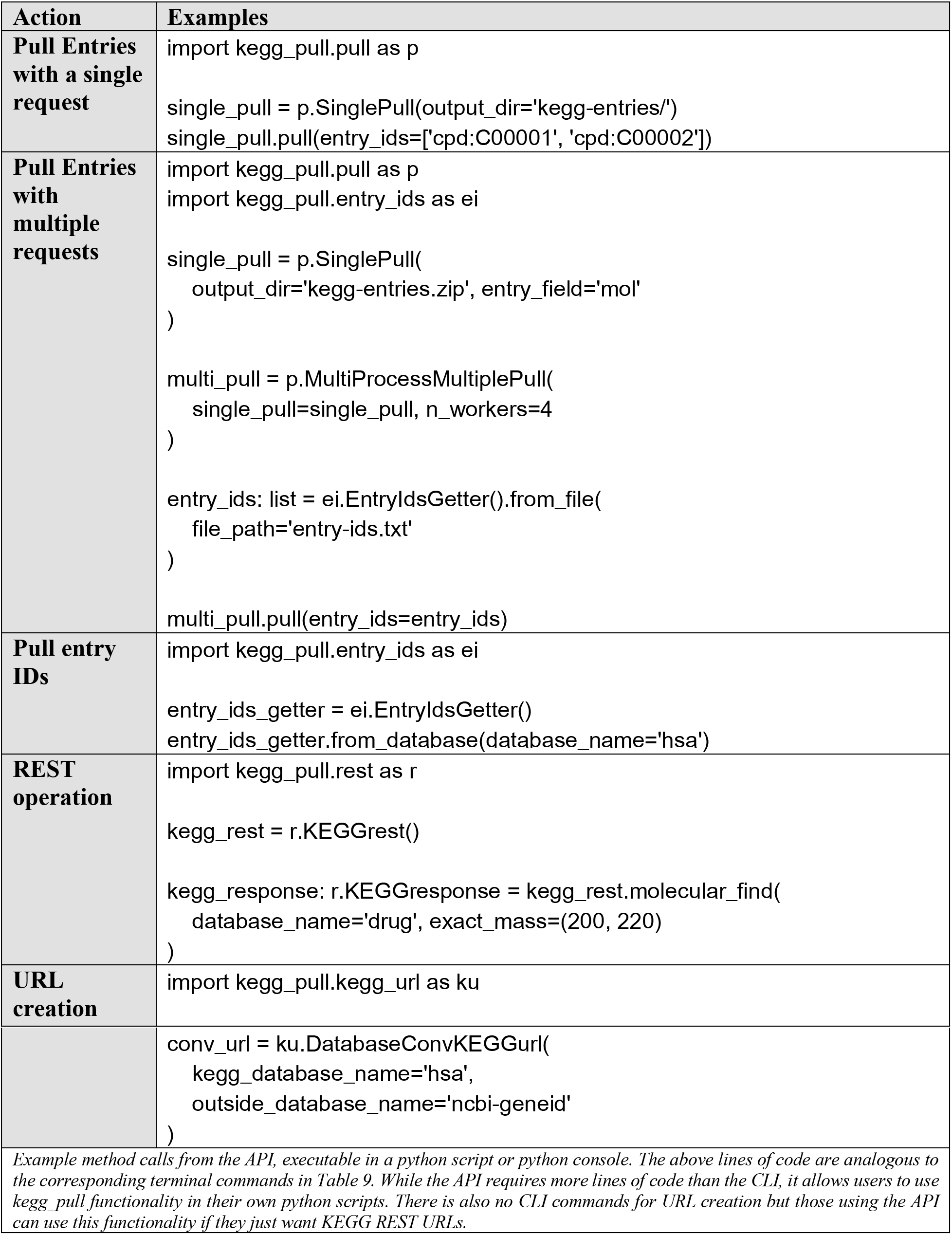
API Examples

| Action | Examples |
| --- | --- |
| **Pull Entries with a single request** | `import kegg_pull.pull as p single_pull = p.SinglePull(output_dir='kegg-entries/') single_pull.pull(entry_ids=['cpd:C00001', 'cpd:C00002'])` |
| **Pull Entries with multiple requests** | `import kegg_pull.pull as p import kegg_pull.entry_ids as ei single_pull = p.SinglePull( output_dir='kegg-entries.zip', entry_field='mol' ) multi_pull = p.MultiProcessMultiplePull( single_pull=single_pull, n_workers=4 ) entry_ids: list = ei.EntryIdsGetter().from_file( file_path='entry-ids.txt' ) multi_pull.pull(entry_ids=entry_ids)` |
| **Pull entry IDs** | `import kegg_pull.entry_ids as ei entry_ids_getter = ei.EntryIdsGetter() entry_ids_getter.from_database(database_name='hsa')` |
| **REST operation** | `import kegg_pull.rest as r kegg_rest = r.KEGGrest() kegg_response: r.KEGGresponse = kegg_rest.molecular_find( database_name='drug', exact_mass=(200, 220) )` |
| **URL creation** | `import kegg_pull.kegg_url as ku` |
|  | `conv_url = ku.DatabaseConvKEGGurl( kegg_database_name='hsa', outside_database_name='ncbi-geneid' )` |
| Example method calls from the API, executable in a python script or python console. The above lines of code are analogous to the corresponding terminal commands in Table 9. While the API requires more lines of code than the CLI, it allows users to use kegg_pull functionality in their own python scripts. There is also no CLI commands for URL creation but those using the API can use this functionality if they just want KEGG REST URLs. |  |

**Table 9.**
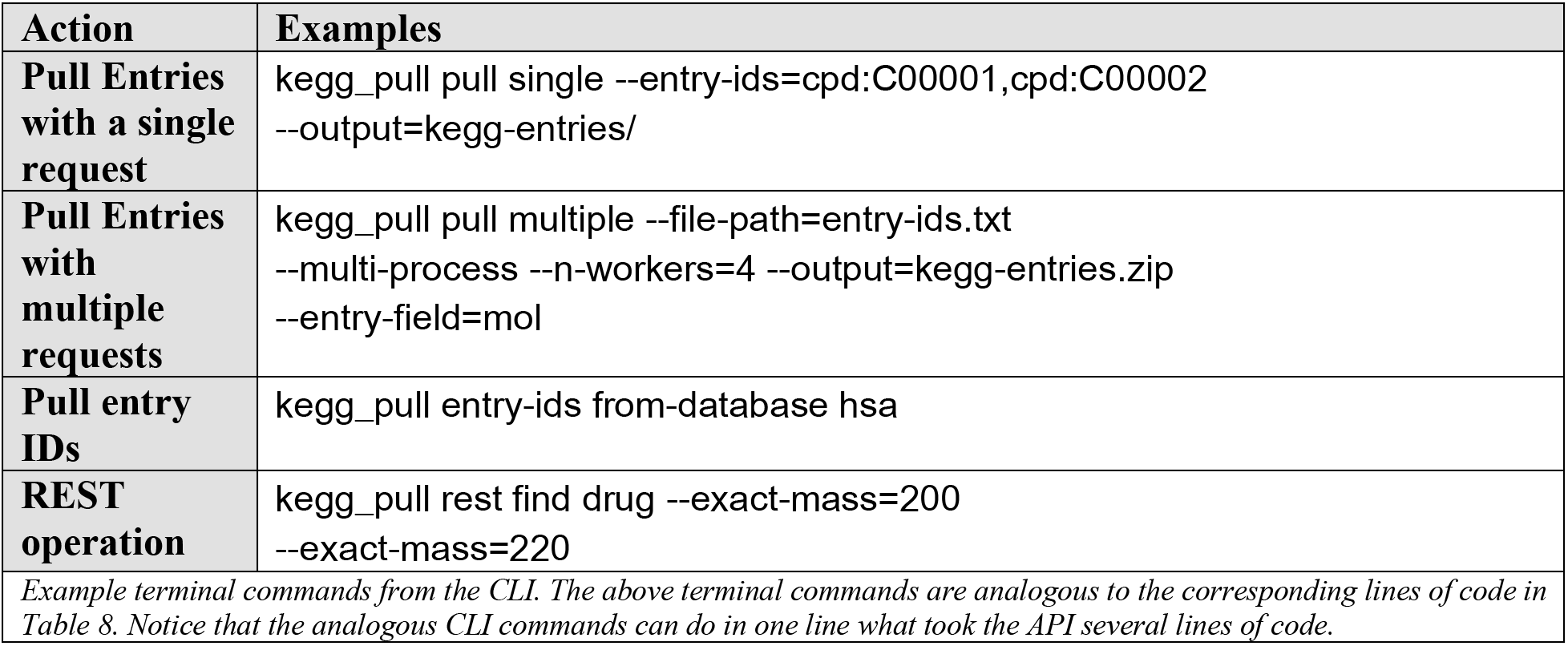
CLI Examples

| Action | Examples |
| --- | --- |
| **Pull Entries with a single request** | kegg_pull pull single --entry-ids=cpd:C00001,cpd:C00002<br>--output=kegg-entries/ |
| **Pull Entries with multiple requests** | kegg_pull pull multiple --file-path=entry-ids.txt<br>--multi-process --n-workers=4 --output=kegg-entries.zip<br>--entry-field=mol |
| **Pull entry IDs** | kegg_pull entry-ids from-database hsa |
| **REST operation** | kegg_pull rest find drug --exact-mass=200<br>--exact-mass=220 |
| Example terminal commands from the CLI. The above terminal commands are analogous to the corresponding lines of code in Table 8. Notice that the analogous CLI commands can do in one line what took the API several lines of code. |  |

## Discussion

Other projects were also considered for the comparison done in Table 10. These projects include KEGG-Crawler with the home page of https://github.com/mentatpsi/KEGG-Crawler, KEGGtools with the home page of https://github.com/FlyPythons/KEGGTools, and django-rest-kegg with the home page of https://pypi.org/project/django-rest-kegg/. They were considered for comparison since they contain code for accessing the KEGG API and downloading KEGG data. However, they give the user no control over which KEGG entries to download but rather choose for the user which entries/data to download, suggesting they are for a more specific purpose than our general purpose kegg_pull package and the other projects compared in Table 10. Additionally, some of these projects are not installable packages but can only be cloned as git repositories, making importing entities into user projects or running scripts on the command line more cumbersome. So we did not deem them appropriate for comparison to kegg_pull.

**Table 10.** Package Information About kegg_pull And Related Packages

| Package Name | Home Page | Python Version | Available Both On GitHub And PyPi | Last Updated |
| --- | --- | --- | --- | --- |
| **kegg_pull** | https://pypi.org/project/kegg-pull/ | >=3.8 | Yes | 2022 |
| **KEGGutils** | https://pypi.org/project/KEGGutils/ | unspecified | Yes | 2020 |
| **biopython (Bio.KEGG.REST)** | https://pypi.org/project/biopython/<br>(https://biopython.org/docs/latest/api/Bio.KEGG.REST.html) | >=3.6 | Yes | 2021 |
| **keggrest** | https://pypi.org/project/keggrest/ | 2.7 | Yes | 2013 |

**Table 11.**
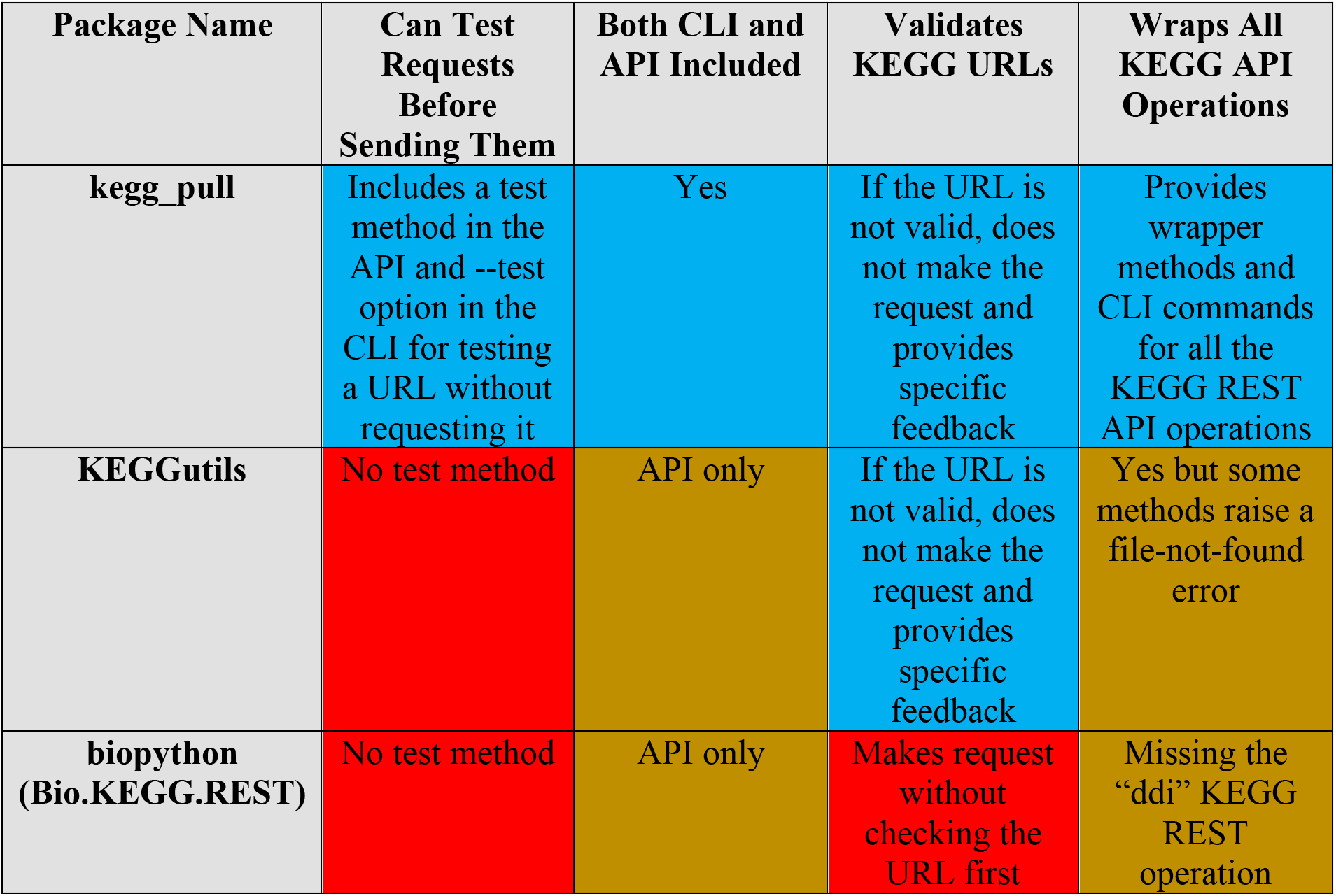
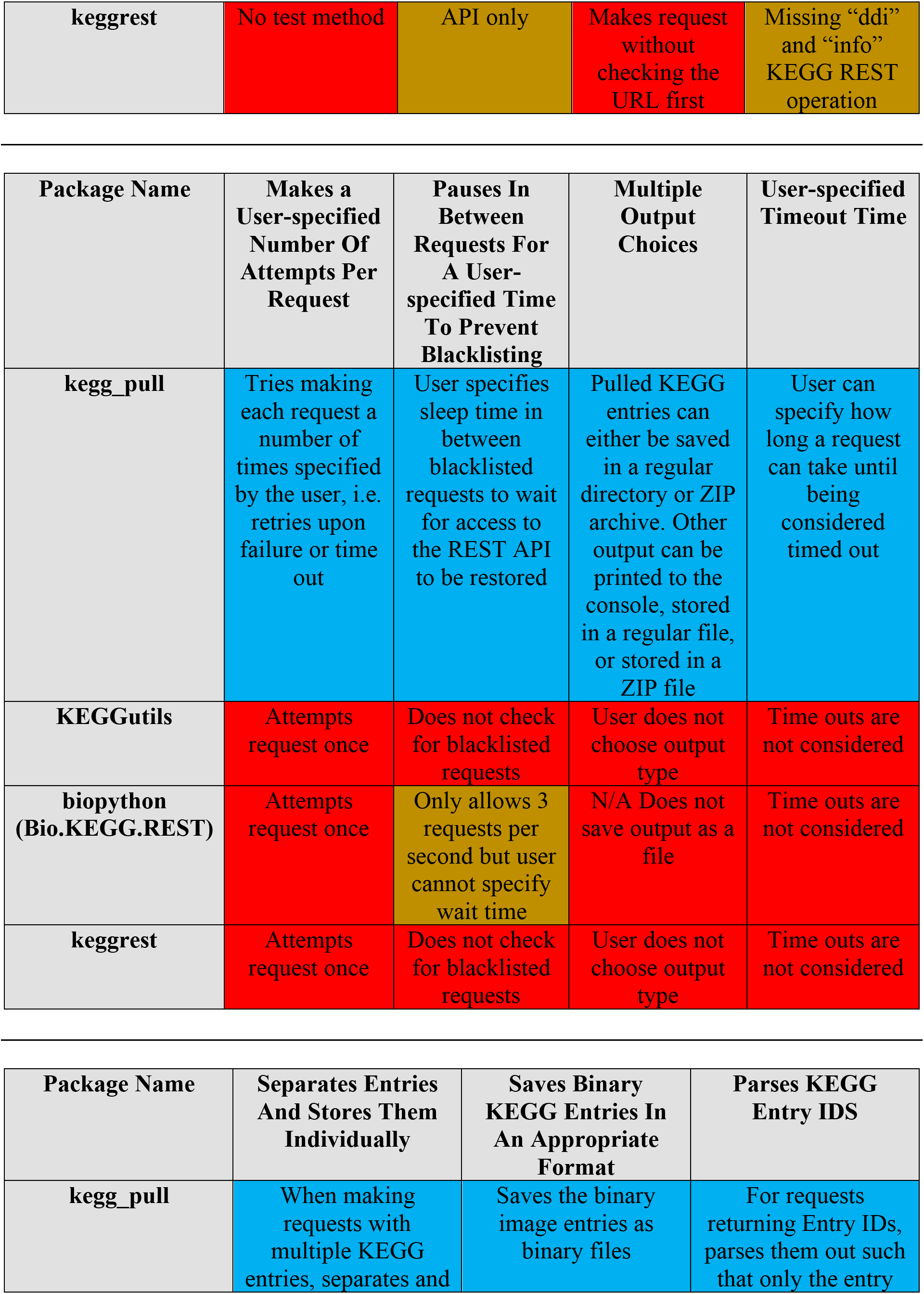
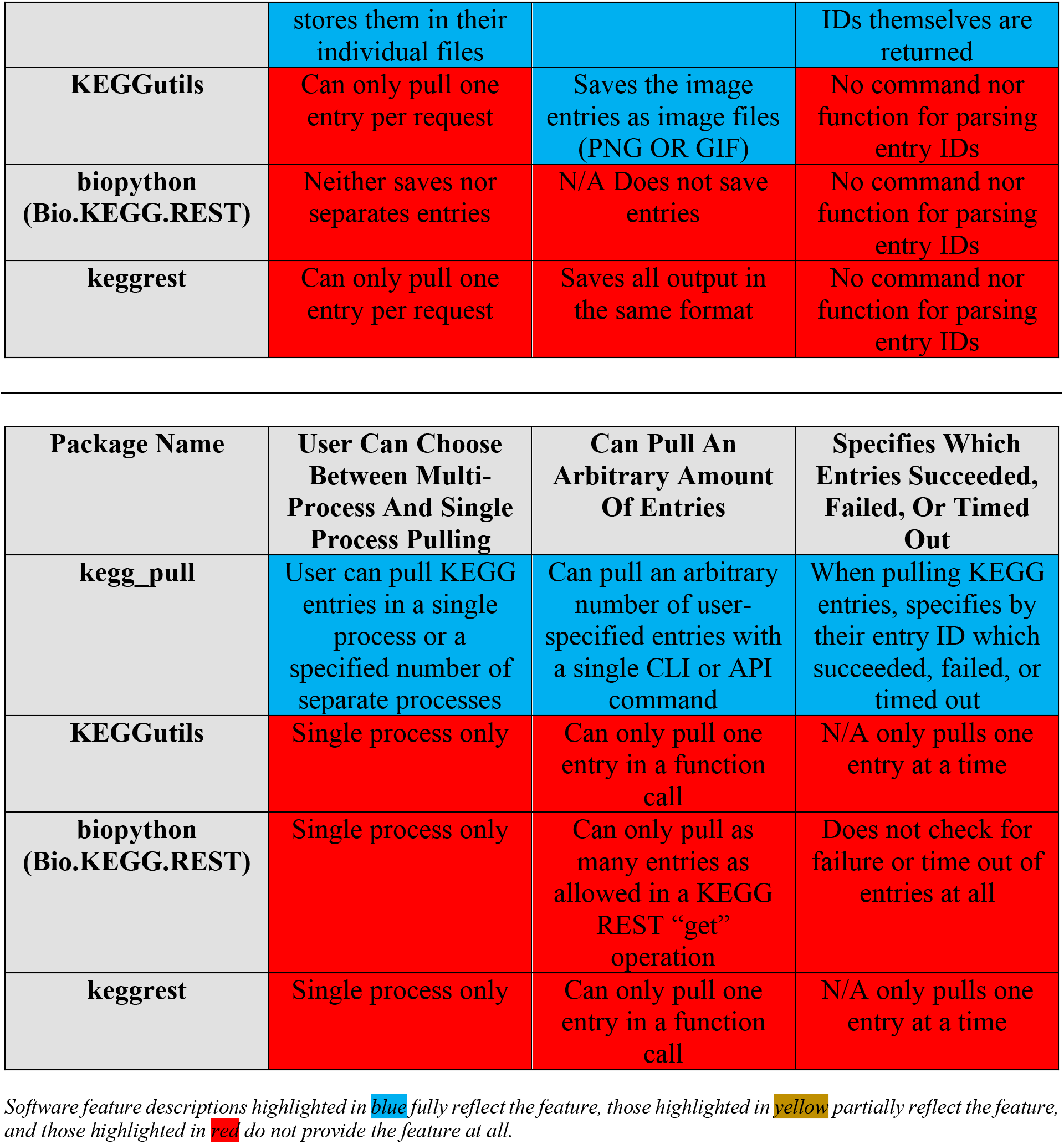
Feature Comparison Of Packages With A similar Purpose As kegg_pull

The new kegg_pull python package makes available the features of the popular R package known as KEGGREST [11] in that it provides an API that wraps the KEGG REST interface, making it easier to make REST requests and doing so in a way that can be automated within user-created Python scripts. While other Python packages (Table 10) [12][13][14] have replicated some of the functionality of KEGGREST, kegg_pull provides a more functional API than all of these packages (Table 11), a complete CLI with a superset of the API functionality (Table 11), and is written to an industrial software engineering standard. Perhaps the most significant feature introduced with kegg_pull is its ability to make multiple requests such that it can pull an arbitrary number of entries with a single command, including the ability to do so in a multi-processing manner. This ability, however, is not without caveats. If a user requests an especially high number of entries in a single call, such as tens of thousands or more, the frequency of blacklisting increases with the number of requested entries. While we cannot prevent blacklisting, the sleep time can be optimized to maximize the success percentage while keeping the overall pull time low. The best sleep time to choose evidently must be higher when requesting a higher number of entries. While there isn’t a mechanism to predict what the best sleep time ought to be ahead of time, we’ve fortunately observed that an overly high sleep time can have negligible effect on the total pull time and pull time can also continue to decrease even after reaching 100 percent success. Therefore, we recommend users lean towards a higher sleep time (e.g. 5.0 or 10.0 s for multiprocessing pulling) as a sleep time that’s too high has negligible effect while still obtaining 100 percent success, but a sleep time that’s too low can both increase the total pull time and lower the success percentage. Extra sleep time is needed when pulling only one entry at a time (e.g. 20.0 s). We recommend that users take advantage of this ability of the KEGG API unless that option is not available for the entries they’d like to pull (i.e. Brite entries and entry fields that don’t support multiple entries within the response body). Considering the increase in success rate when pulling multiple entries per request as well as the significant decrease in pull time, it could be helpful for both users of kegg_pull and users of the KEGG API in general if KEGG both enabled support for pulling multiple entries for all entry and entry field types and even allowing more than 10 entries to be requested. All this applies to multi-processing, whereas the sleep time is not as important in single processing. As we’ve seen, even a sleep time of 0.0 can result in 100 percent success, likely because the time in between requests is already necessarily higher, preventing black listing.

Since it’s still possible for entry requests to fail, we recommend users re-run kegg_pull on the failed entries after doing their best to initially select a good sleep time. This is not just because of blacklisting, but entries can inadvertently fail for other reasons such that they may succeed the second time. Entries that continuously fail to be pulled may be considered no longer available, as with the 20 consistently failed Brite entries. In such cases generally and in the case of the Brite database specifically, we recommend that KEGG either remove the IDs of these available entries from the output of the “list” REST operation or that they troubleshoot to see whether these entries can be made available.

We recommend the --force-single-entry flag (CLI) or force_single_entry parameter (API) to be set if brite entry IDs are included in the call. While if a user chooses to pull the entire brite database, kegg_pull is smart enough to only pull one entry at a time. But it can’t know to do this if a file containing KEGG entries is provided.

Pulling KEGG entries into a ZIP archive is significantly slower both when multi-processing and when single processing. Likewise, single process pulling is significantly slower than multi-process pulling, both when pulling into a ZIP archive and when pulling into a regular directory. This means that multi-processing is still worth performing for ZIP archives despite locking multi-process unsafe code. Table 12 specifies the best decisions between multi-processing versus single processing and ZIP archives versus regular directories depending on the circumstances.

**Table 12.** Recommendation For Multi-processing And Storage Options

| Circumstance | Multiple Cores<br>Available | Only One Core<br>Available |
| --- | --- | --- |
| **Must Store In ZIP**<br>**Archive** | Multi-processing pull into<br>a ZIP archive | Single processing pull<br>into a ZIP Archive |
| **No Need For ZIP**<br>**Archive** | Multi-processing pull into<br>a regular directory | Single processing pull<br>into a regular directory |
| *The recommended manner of pulling based on the circumstance, with the need for a ZIP archive on the left side and the amount of cores available on the top.* |  |  |

## Conclusions

The kegg_pull Python package provides the richest programmatic and command line access to the KEGG API to date. The clean objected oriented implementation provides robust multiprocessing KEGG entry retrieval (pull) functionality that can be used in user-created Python scripts or in data analysis pipelines and workflow managers, thus improving the FAIRness of KEGG. The package is implemented to a high industrial software engineering standard, which includes both unit and integration tests that provides 100% code coverage. The code base is revision controlled and managed on GitHub, documentation is auto-updated onto associated GitHub Pages, and the package is distributed through the Python Package Index. Feedback is greatly appreciated. Any potential bugs or requests for new features can be submitted on our GitHub repository issues page here:

https://github.com/MoseleyBioinformaticsLab/kegg_pull/issues.

## Availability And Requirements

**Project name:** kegg_pull

**Project home page: https://github.com/MoseleyBioinformaticsLab/kegg_pull**

**Operating system(s):** Platform independent

**Programming language:** Python

**Other requirements:** Python3.8 or higher

**License:** Modified BSD 3 License

**Report Bugs And Feature Requests Here:**

https://github.com/MoseleyBioinformaticsLab/kegg_pull/issues

## List Of Abbreviations

API: Application programming interface
CLI: Command line interface
KEGG: Kyoto Encyclopedia of Genes and Genomes
REST: Representational State Transfer
URL: Uniform resource locator

## Declarations

### Ethics Approval And Consent To Participate

Not Applicable

### Consent For Publication

Not Applicable

### Availability Of Data And Materials

GitHub repository: https://github.com/MoseleyBioinformaticsLab/kegg_pull

Python Package Index (PyPi): https://pypi.org/project/kegg-pull/

Documentation: https://moseleybioinformaticslab.github.io/kegg_pull/

Figshare containing this manuscript’s table results and the scripts to produce them: https://doi.org/10.6084/m9.figshare.21471990

### Authors Contributions

EH and HNBM created the objected oriented design in multiple prototype-redesign cycles. EH implemented the software, automated unit and integrative testing, automated end-user documentation generation, and automated package distribution via PyPI. EH wrote the package documentation. HNBM reviewed both the implementation and package documentation. EH wrote the initial draft of the manuscript. HNBM and EH revised the manuscript in multiple revision rounds. HNBM provided support via funded grants.

## Acknowledgements

Not Applicable

## Supplementary Material for

### Brite Failed Entries

br:br03220

br:br03222

br:br01610

br:br01611

br:br01612

br:br01613

br:br01601

br:br01602

br:br01600

br:br01620

br:br01553

br:br01554

br:br01556

br:br01555

br:br01557

br:br01800

br:br01810

br:br08011

br:br08020

br:br08120

### Single Process Pull Success Percentage And Time Spent Pulling By Sleep Time (KO Database)

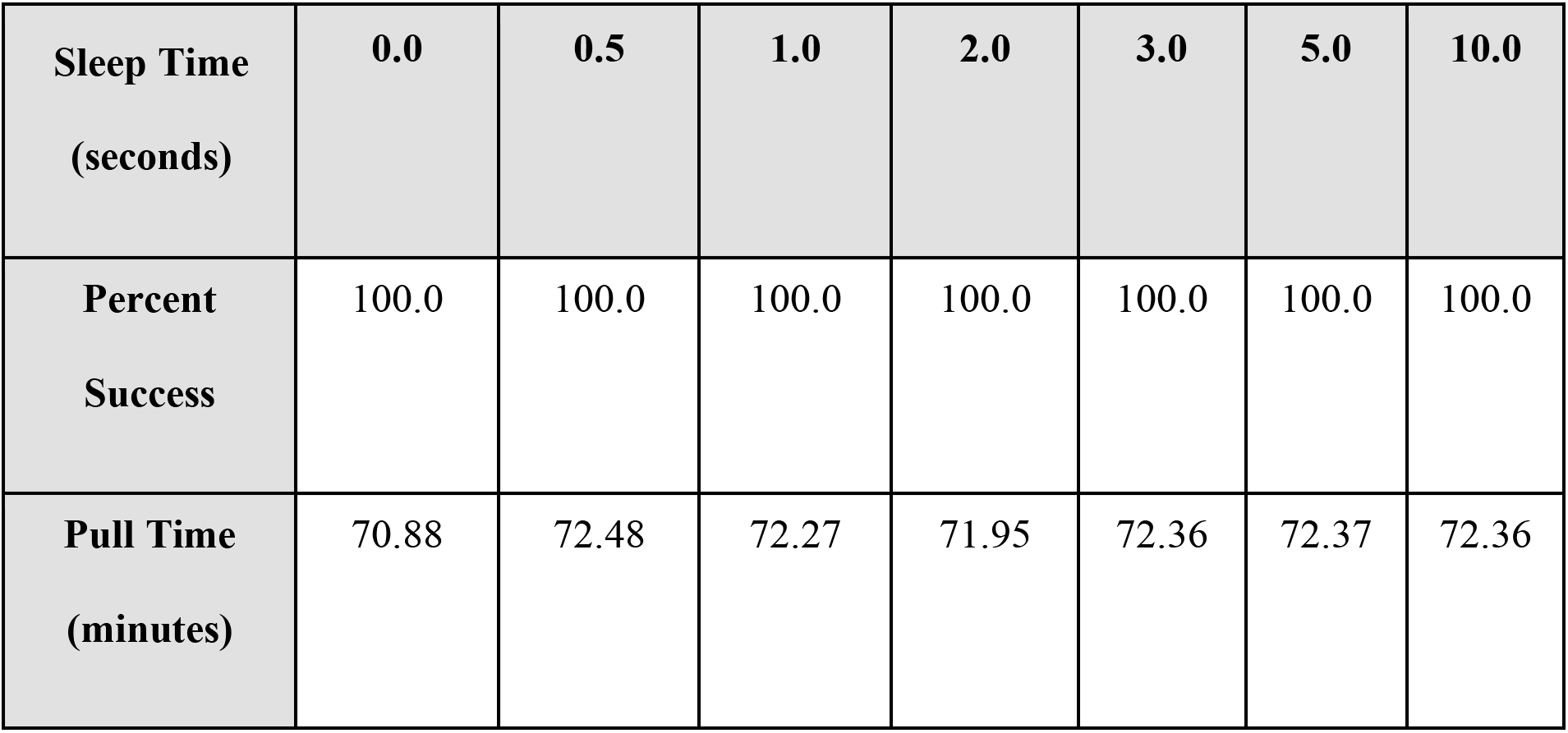

| **Sleep Time**<br>**(seconds)** | **0.0** | **0.5** | **1.0** | **2.0** | **3.0** | **5.0** | **10.0** |
| --- | --- | --- | --- | --- | --- | --- | --- |
| **Percent**<br>**Success** | 100.0 | 100.0 | 100.0 | 100.0 | 100.0 | 100.0 | 100.0 |
| **Pull Time**<br>**(minutes)** | 70.88 | 72.48 | 72.27 | 71.95 | 72.36 | 72.37 | 72.36 |

